# Estrogen Receptor Expression Changes After Puberty in the Porcine Anterior Cruciate Ligament

**DOI:** 10.64898/2026.03.09.710593

**Authors:** Jacob D. Thompson, Matthew B. Fisher

**Author notes:** Corresponding Author Contact: Address: 4130 Engineering Building III, 911 Oval Drive, CB 7115, Raleigh, NC, 27695. **Author Contributions:** J.D.T and M.B.F. [1] made substantial contributions to research design, acquisition, analysis, and interpretation of data; [2] drafted the paper and revised it critically; and [3] approved of the submitted and final versions.

## Abstract

Anterior cruciate ligament (ACL) injuries disproportionately affect female adolescent athletes, with hormonal influences implicated in this sex disparity. However, the relationship between pubertal hormonal changes and ACL gene and protein expression remains poorly understood. This study characterized hormone receptor expression and transcriptional profiles in the anteromedial (AM) and posterolateral (PL) bundles of female porcine ACLs before and after puberty. ACL bundles were collected from pre-pubescent (8 weeks) and post-pubescent (>8 months) female Yorkshire cross-breed pigs (n=6/group) and analyzed using gene expression profiling, western blotting, and immunofluorescence. Pre-pubescent ACLs exhibited greater expression of primary matrix genes (COL1A1, COL1A2, ELN, TNMD), suggesting active matrix synthesis, while post-pubescent ACLs showed elevated secondary matrix genes (COL3A1, LUM, COMP), indicating a homeostatic state. Notably, estrogen receptor alpha (ERα) gene and protein expression were significantly greater in post-pubescent ACLs, particularly in AM bundles, whereas G-protein coupled estrogen receptor (GPR30) expression was elevated pre-puberty. Both receptors were distributed homogeneously throughout the tissue. Progesterone receptor protein expression was not detected in any samples. Histologically, post-pubescent ACLs demonstrated decreased cellularity and thicker fascicles compared to pre-pubescent tissues. These findings indicate that ACL sensitivity to estrogen varies across development, with increased ERα expression post-puberty potentially rendering the ligament more responsive to circulating estrogen. This work provides foundational evidence for age-dependent hormonal responsiveness in the ACL and motivates further investigation into how sex hormones influence ACL injury risk in adolescent females.

## Introduction

Musculoskeletal injuries continue to pose a significant burden on society, with an estimated economic cost of $381 billion annually in the United States ^1^. Among these injuries, anterior cruciate ligament (ACL) ruptures are particularly prevalent in young athletes ^2–4^. The increasing trend of early sports specialization and heightened competition in youth sports has contributed to a rise in pediatric ACL injuries ^5–8^. Furthermore, female adolescent athletes are at a 3-4 times higher risk of tearing their ACL within specific sports settings, like basketball or soccer ^9–13^. This sex disparity in injury rates has been partly attributed to hormonal influences. Early studies have identified associations between menstrual cycle phases and ACL injury timing, particularly during the follicular phase near ovulation when estrogen spikes, implicating estrogen in heightened injury susceptibility among females ^14–17^. In humans where hormone levels were tracked, anterior knee joint laxity has also been shown to increase around 1 – 2 mm in women three days following the estradiol spike immediately preceding ovulation ^18^. However, in animal studies, there have been inconsistent findings about the impact of sex hormones on ACL mechanics, likely from variability in approaches and differences in experimental timeframes ^19–22^. Despite the previous work performed, how physiological changes in puberty relate to changes in the ACL and subsequent injury risk is still unclear.

Sex hormone receptors have been identified within the ACL in humans and rodents for estrogen, progesterone, and testosterone ^23,24^. Sex hormone receptors, including nuclear estrogen receptors alpha (ERα) and beta (ERβ), g-coupled protein estrogen receptor (GPR30), nuclear progesterone receptors alpha (PRα) and beta (PRβ), and androgen receptors (AR) are all critical components for how sex hormones enact transcriptional changes in cells. Direct cell responses to sex hormones have been explored, and have shown that ACL fibroblasts are sensitive to both estradiol and progesterone, with decreases in proliferation, type I collagen synthesis, and collagen crosslinking through lysyl oxidase from estradiol exposure, and an increase in proliferation and collagen synthesis from progesterone exposure ^25–27^. There are no studies, however, characterizing how hormone receptors might change throughout growth. Such data can provide more insights into when and how the ACL might respond to hormonal fluctuation, but the ability to obtain uninjured ACL tissue in skeletally immature human subjects is extremely challenging.

Animal models can overcome this limitation, and the adolescent pig model has been shown to be an effective model in characterizing structural and mechanical changes in the ACL throughout skeletal maturation ^28–30^. Sex-specific differences have also been described in the adolescent pig where the anteromedial (AM) bundle size increases throughout growth for males and females, but the posterolateral (PL) bundle in females plateau around mid-adolescence ^31^. The AM bundle of the male ACL also becomes more dominant than the PL bundle at an earlier point than females ^31^. The mechanisms, however, driving these bundle-specific changes are still unknown. Pigs also undergo an estrus cycle that begins with the female pigs going into their first heat. Prior to puberty in pigs, hormone levels of estradiol and progesterone are low or not-detectable, but after puberty, pigs undergo a multi-week cycle time (21 days) with dynamic estrogen and progesterone peaks ^32^. For these reasons, the pig model is used to study reproductive health ^33,34^. This motivates the pig model as a promising model to better understand biological changes in the ACL and its bundles in the context of the changing hormonal environment across growth. Therefore, the objective of this study was to characterize transcriptional changes and receptor protein differences between the AM and PL bundle in pre- and post-pubescent female pigs and understand more specifically how hormone receptors in the ACL differ. We hypothesized that there would be distinct transcriptional differences before and after puberty and greater hormone receptor expression in post-pubescent female pig ACLs compared to pre-pubescent female pig ACLs. We also hypothesized that there would be minimal differences between bundles at each age. We tested these hypotheses by collecting ACL bundles from female pigs at pre-pubertal and post-pubertal ages, extracting RNA for gene expression, generating protein lysates for hormone receptor expression levels, and staining the ACL bundles for hormone receptors.

## Methods

### Specimen Collection

The AM and PL bundles of the ACL were collected from female pigs (Yorkshire cross-breed) prior to puberty (approximately 8 weeks old) or post-puberty (>8 months old) (n=6 animals/age; Figure 1). Puberty in Yorkshire pigs occurs between 5 and 7 months of age ^35^. All tissues were collected in the morning from animals post-euthanasia (sodium pentobarbitol solution) from separate studies performed at North Carolina State University and approved by IACUC. A sample size of 6 animals was determined based on availability of post-pubescent female pigs. The bundles were immediately snap-frozen in liquid nitrogen for gene expression and western blot analysis or placed in optimal cutting temperature for immunofluorescence and histological staining for primary experimental outcomes. Uterine tissue was also harvested as a positive control for hormone receptors. On a subset of the post-pubescent animals collected (n=3), ovaries were harvested as a secondary outcome to verify sexual maturity and serum was collected prior to euthanasia and stored at -80°C until further use.

**Figure 1.**
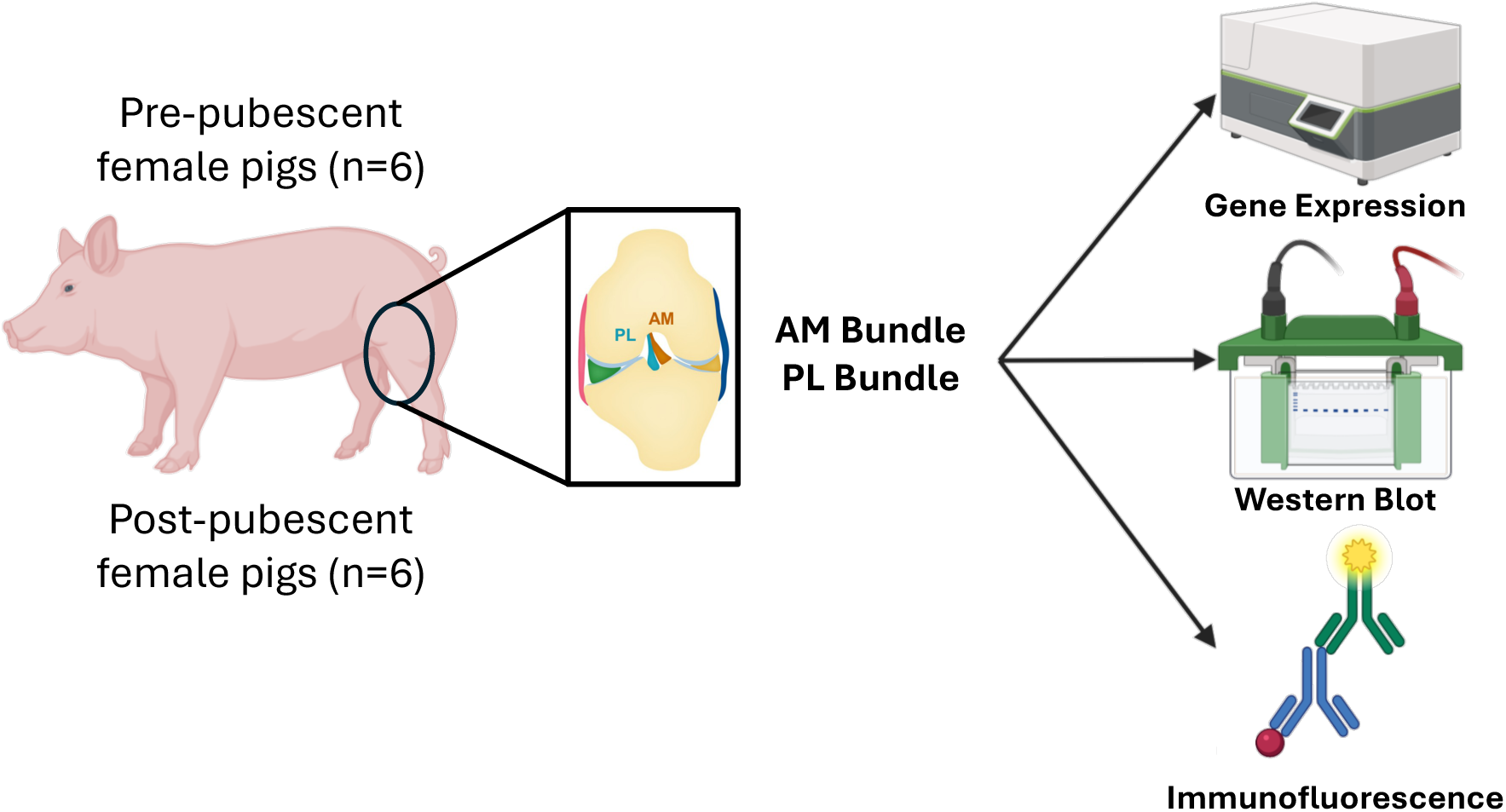
Schematic highlighting study design and approach to characterize genetic differences and hormone receptor expression differences between the AM and PL bundles across growth. Pre-and post-pubescent female pigs had their ACLs removed, separated by bundle, then characterized using gene expression, western blot, and immunofluorescence. Created in BioRender. Fisher, M. (2026) https://BioRender.com/yjybb9p

### Serum Hormone Quantification

Serum was collected on a subset of animals prior to euthanasia, and these samples were transported to the Clinical Endocrinology Laboratory at the University of California, Davis for hormone analysis, who were blinded to the age and reproductive status of each animal. Serum estradiol was quantified using the ultra-sensitive estradiol radioimmunoassay kit (DSL-4800, Beckman Coulter, Brea, CA, USA), and serum progesterone concentrations were measured using enzyme-linked immunosorbent assays (ELISAs) developed and validated in-house. The RIA for estradiol exhibited an intra-assay coefficient of variation (CV) of 8.4%, while the intra-assay CV for the progesterone ELISA was 8.6%.

### Gene Expression Analysis

Total RNA from the ACL bundles of pre- and post-pubescent female animals was harvested with an ENZA total RNA kit (Omega Bio-tek Inc., Norcross, GA, USA) following TRIzol and chloroform extraction in a Bead Mill Homogenizer (ThermoFisher Scientific, Waltham, MA, USA). RNA yield and quality were assessed by UV spectroscopy (NanoDrop 2000, ThermoFisher Scientific, Waltham, MA, USA) where a minimum of 100 ng of RNA was required for gene expression analysis. RNA purity was determined by the A_260_ / A_280_ ratio with purities > 1.7. Transcriptional analysis was performed with the nCounter MAX Analysis System and data was processed using nSolver 4.0 (NanoString Technologies, Bruker Spatial Biology Inc., Seattle, WA, USA). A custom code-set of 45 genes was used to analyze extracellular matrix protein-related genes and hormone receptor genes, including *ESR1* (encodes estrogen receptor alpha), *ESR2* (encodes estrogen receptor beta), *GPER1* (encodes g-coupled estrogen receptor), and *PGR* (encodes progesterone receptor) (Table S1). Gene expression data was normalized to the positive control reference genes included within the custom code set (*HPRT1*, *GAPDH*, and *ACTB*), and the average of the negative controls was used to background subtract all counts.

### Western Blot

Total protein from the ACL bundles of pre- and post-pubescent female pigs was isolated with RIPA buffer and 1X HALT protease inhibitor cocktail with a Bead Mill Homogenizer (ThermoFisher Scientific, Waltham, MA, USA). Homogenized samples were spun at 16,000xg for 20 minutes at 4°C and the supernatant was collected. Total protein concentration was determined by a Pierce BCA Protein assay kit (ThermoFisher Scientific, Waltham, MA, USA). Protein lysates were reduced and boiled at 95°C then separated by size on 4-12% NuPage gels as per the manufacturer’s instructions. The protein bands were transferred to 0.45 µm PVDF membranes and blocked with 5% milk. They were then blotted for estrogen receptor alpha (1:1000 ERα, PA1-309 ThermoFisher), g-coupled protein estrogen receptor (1:800 GPR30, PA5-28647 ThermoFisher), and GAPDH (1:1000, 2118S Cell Signaling Technologies) overnight in 5% BSA at 4°C. Progesterone receptor beta (PRβ, 3157 Cell Signaling Technologies) could not be validated from the positive uterine control in western blot, so was excluded from this analysis. A secondary antibody was used to visualize hormone receptor bands (anti-rabbit 1:20000, A32735 ThermoFisher) in 5% milk + 0.01% SDS for 2 hours at room temperature. Blots were imaged with a LI-COR Odyssey CLx imager (LI-COR, Lincoln, NE, USA), and relative protein levels were quantified in ImageJ after normalization to GAPDH. A uterus control was run on each gel as a positive tissue control for hormone receptors and to account for different signal intensities across gels.

### Immunofluorescence and Histological Analysis

ACL bundles for pre- and post-pubescent female pigs were cryosectioned at 10 µm. For immunofluorescence, sections were permeabilized for 15 minutes in 0.1% Triton-X in 1X PBS and blocked in a solution of commercial IHC/ICC blocking buffer (45%), 0.1% Triton-X (45%), and donkey serum (5%) for 1 hour at room temperature. Primary antibodies were then added for ERα (1:400), PRβ (1:200), and GPR30 (1:500) in 1% bovine serum albumin, 0.3% Triton-X in 1X PBS overnight at 4°C. Rabbit IgG isotype, no primary, and positive uterus tissue controls were used to determine the non-specific binding of the primary and secondary antibodies and to verify positive staining (Figure S3). Secondary antibodies were added for 2 hours at room temperature in the dark and slides were mounted with Prolong Gold with DAPI. All sections were imaged with a cellSens Olympus LS fluorescent microscope (Olympus, EvidentScientific, Waltham, MA, USA). Cellularity was determined by the analyze particles function within ImageJ.

Cryosections for histological analysis were brought to room temperature and fixed on the slide with 4% paraformaldehyde for 15 minutes. They were then washed with 1X PBS and stained with picrosirius red, hematoxylin & eosin, and alcian blue to assess matrix organization across age and bundle.

### Statistical Analysis

Nanostring gene expression raw counts for all samples were normalized by housekeeping controls and built-in negative controls. They were utilized to assess the top 10 expressed genes for each group. The log2 of normalized count data was used in the remainder of the work to determine differences between age and bundle. Comparisons between bundle and age were made using multiple t-tests. Multiple paired t-tests were used when comparing AM and PL bundles, and multiple unpaired t-tests were used when comparing pre- and post-pubescent ACL tissues. Multiple comparisons were controlled by Holm-Sidak post-hoc tests. For western blots, relative band density was normalized to GAPDH for each well and normalized across gels by the matched uterine control. One outlier was removed from western blot analysis via ROUT analysis with Q=1%, and a two-way ANOVA with paired AM and PL bundles was used to determine the effect of age and bundle. Multiple comparisons were controlled by Sidak post-hoc tests. All statistical analyses were run with a set alpha value of 0.05.

## Results

### Verification of Sexual Maturity in Post-Pubescent Animals

A subset of the post-pubescent animals (3/6 animals analyzed based on availability) used for this study had serum collected and analyzed for concentrations of estradiol and progesterone at the time of euthanasia. The estradiol values ranged from 5.35 to 49.24 pg/ml, and progesterone ranged from 0.49 to 3.77 ng/ml (Figure 2A). Post-pubescent animals also had their ovaries removed and observed to verify cycling status (Figure 2B). All the ovaries had both follicles and corpora lutea present, indicating an active estrus cycle. However, specimens one and two had more apparent transparent follicles, which indicates they are closer to estrus, compared to specimen three that had darker, translucent follicles.

**Figure 2.**
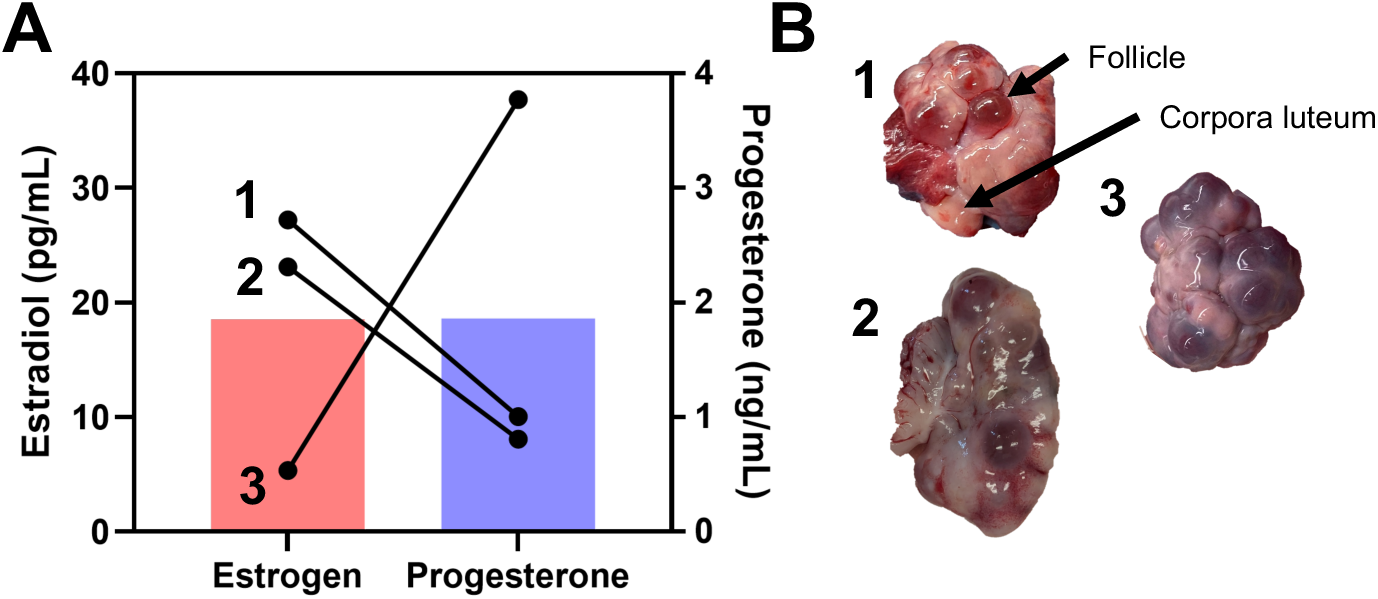
Verification of sexual maturity and cycle phase in post-pubescent animals. (A) Serum estradiol and progesterone values from three of the six animals and (B) images of ovaries from three of the six animals to demonstrate active cycling in the post-pubescent group. Both follicles and corpora lutea are present within all ovaries collected.

### Transcriptional Shifts in the ACL Across Age

Transcriptional differences were assessed in the AM and PL bundles from pre- and post-pubescent animals (6/6 pre-pubescent and 6/6 post-pubescent animals analyzed, Figure 3A). Strong expression of extracellular matrix genes (collagen 1α1, collagen 1α2, decorin, collagen 3α1, collagen 6α1, and lubricin) was found across both age and bundle (Figure S1). Other highly expressed genes include proteoglycans (fibromodulin, biglycan, lumican) and glycoproteins (fibronectin and tenomodulin) relevant to ligament formation. There were several differentially expressed genes when directly comparing across age in the AM bundle (Figure 3B, Figure S2). Bone-morphogenic protein 12 (*GDF7/BMP12*), lumican (*LUM*), collagen 3α1 (*COL3A1*), and estrogen receptor α (*ESR1*) all had higher expression in post-pubescent AM bundles compared to pre-pubescent AM bundles. In the pre-pubescent AM bundle, there was greater expression of collagen 1α1 (*COL1A1*), collagen 1α2 (*COL1A2*), collagen 2α1 (*COL2A1*), elastin (*ELN*), collagen 6α1 (COL6A1), transient receptor potential cation channel subfamily V member 4 (*TRPV4*), tenomodulin (*TNMD*), and g-coupled protein estrogen receptor (*GPER1*) compared to the post-pubescent AM bundle. When comparing PL bundles across age, there were only two differentially expressed genes (Figure 3C). Cartilage oligomeric protein (*COMP*) had greater expression in post-pubescent PL bundles, and *TNMD* had greater expression in pre-pubescent PL bundles. Across bundles, there were little differences in gene expression within either age group. In pre-pubescent animals, the PL bundle had higher expression of a disintegrin and metalloproteinase with thrombospondin motifs 4 (*ADAMTS4*) (Figure 3D). There were no differences between the AM and PL bundles in the post-pubescent animals (Figure 3E).

**Figure 3.**
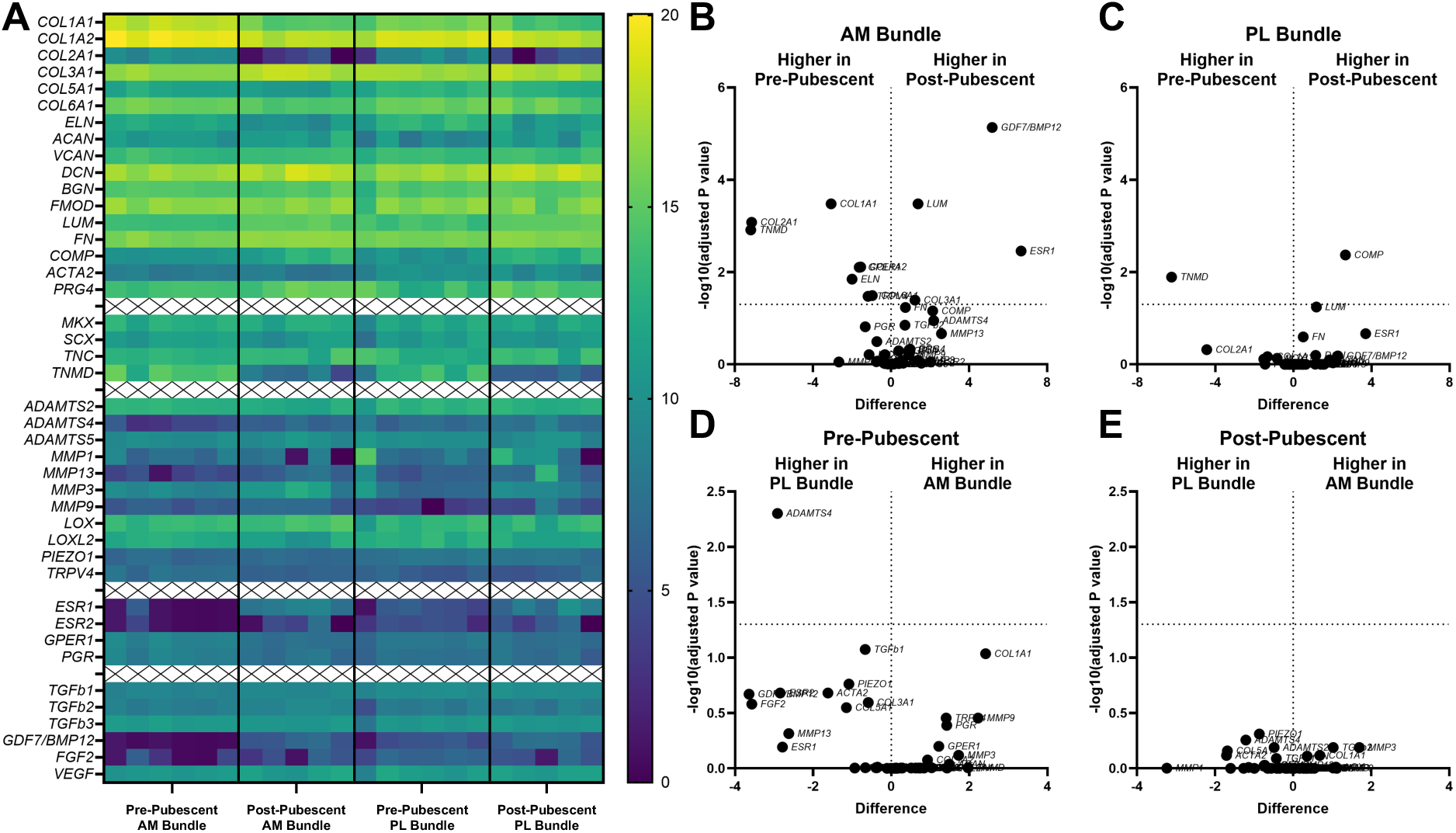
Gene transcription varied before and after puberty, but less so between AM and PL bundles. (A) Heat map showing log2 of all gene counts for AM and PL bundles across development. Differentially expressed genes between pre- and post-pubescent (B) AM and (C) PL bundles. Differentially expressed genes between AM and PL bundles in (D) pre-pubescent and (E) post-pubescent animals. Tissues from individual animals shown within heat map and volcano plots of log2 sample differences and -log10 adjusted p-values from multiple comparisons.

### Greater Expression of Estrogen Receptor in Post-Pubescent AM Bundles

Looking more specifically at hormone receptor gene expression, *ESR1* and *GPER* had different expression levels in the post-pubescent AM bundles compared to pre-pubescent AM bundles (adj p=0.003, adj p=0.008, respectively; Figure 4A,B). These differences were also opposing, where *ESR1* was significantly greater in post-pubescent AM bundles, whereas *GPER1* was significantly greater in pre-pubescent AM bundles. *ESR1* was also greater in the post-pubescent PL bundles compared to pre-pubescent PL bundles, but this was not statistically significant (adj p=0.22). *GPER1* had similar expression levels for the PL bundles across. *ESR2* had high variability and no statistically significant differences across groups (Figure 4C). *PGR* had expression levels similar to that of *GPER1*, with the highest expression in the pre-pubescent AM bundle, but there were no statistically significant differences across any groups (Figure 4D).

**Figure 4.**
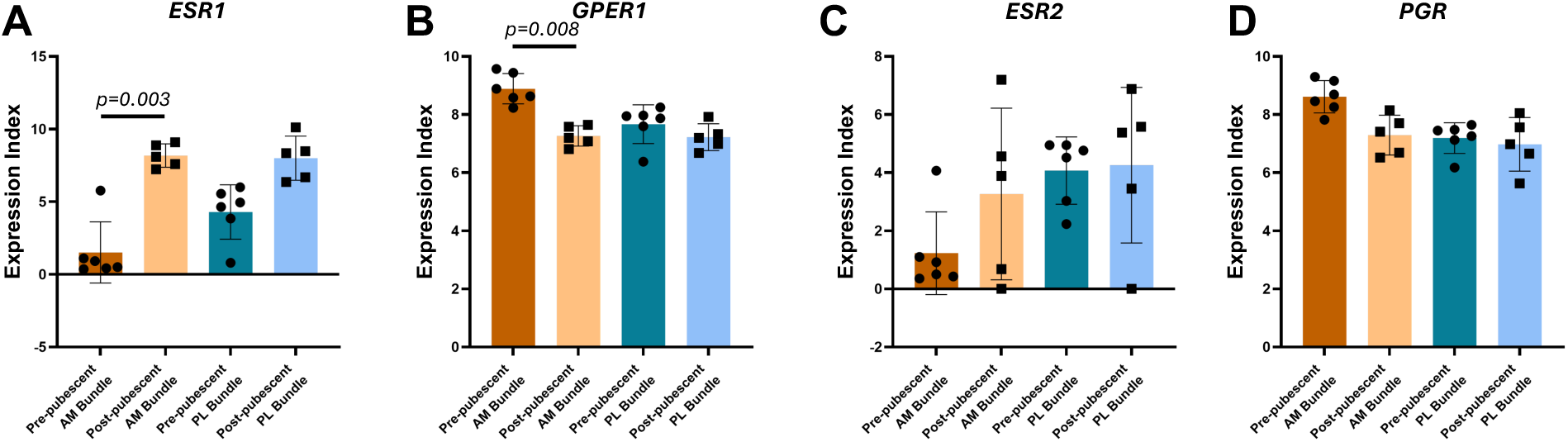
Estrogen receptor alpha and g-coupled protein estrogen receptor gene expression varies between pre- and post-pubescent ACLs. (A) Estrogen receptor alpha (*ESR1*), (B) g-coupled protein estrogen receptor (*GPER1*), (C) estrogen receptor beta (*ESR2*), and (D) Progesterone receptor (*PGR*) expression between groups. Each data point is a biological replicate. Data expressed as mean ± s.d of expression index (log2 of RNA transcript counts). Bars indicate statistical significance between pre- and post-pubescent samples adjusted for multiple comparisons.

### Protein Expression of ERα Greater in Post-Pubescent ACLs

Given gene expression differences in *ESR1* and *GPER1*, protein quantification for these receptors was carried out through western blots (6/6 pre-pubescent and 6/6 post-pubescent animals analyzed, Figure 5A,B, Figure S3). The ERα protein (encoded by *ESR1*) was found primarily in the post-pubescent ACLs, with minimal to no expression in the pre-pubescent group aside from one animal. Comparing across age, there was a significantly greater amount of ERα protein in the post-pubescent AM bundles and PL bundles compared to the pre-pubescent AM and PL bundles (Figure 5C; adj p=0.01, adj p=0.01 respectively). Outside of the expected molecular weight of approximately 66 kDa, there were other non-specific bands seen in both pre and post-pubescent samples for the ACL (Figure S3A). GPR30 (encoded by *GPER1*) protein was found within all specimens tested, but the expression was not significantly different across age or bundle (Figure 5D). However, on average, there was slightly greater GPR30 protein expression in the pre-pubescent AM bundle compared to the post-pubescent AM bundle (adj p=0.11). Additional bands representing glycosylated, modified, or bound forms of GPR30 or non-specific binding were seen in all samples (Figure S3B).

**Figure 5.**
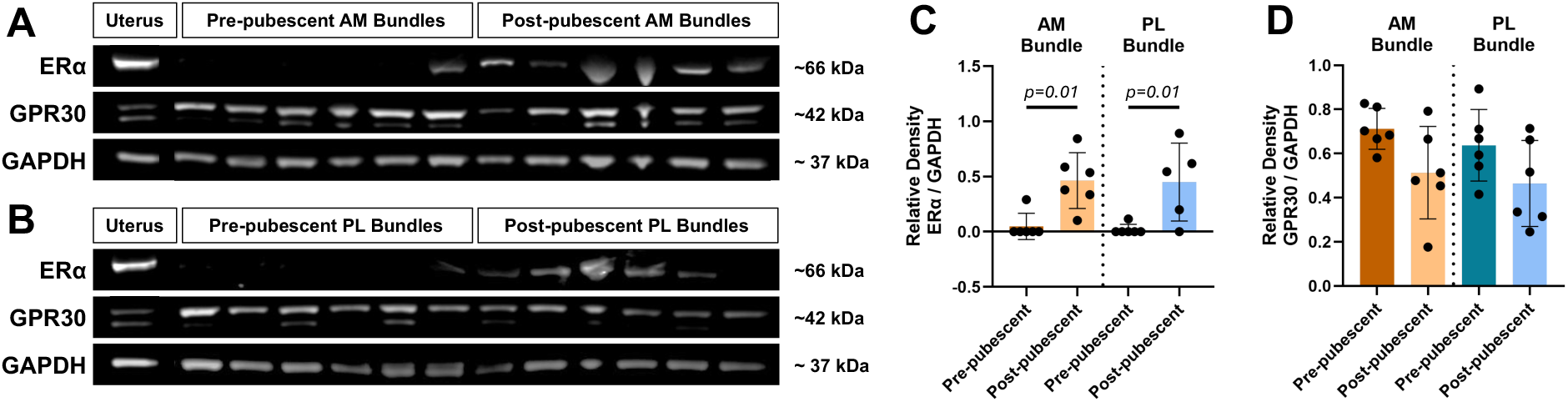
Estrogen receptor protein expression changes with age and varies between animals. Estrogen receptor protein expression from western blots of pre- and post-pubescent animal (A) AM and (B) PL bundles. Quantification of relative band density for (C) ERα across age and bundle and (D) GPR30 across age and bundle. Each data point is a biological replicate. Data expressed as mean ± s.d. Bars indicate statistical significance.

### Similar Spatial Distribution of ERα Across Age

Spatially, positive signal for both ERα and GPR30 was observed throughout the tissue for both the AM bundle (Figure 6) and PL bundle (Figure S5). Focusing on the AM bundle, both estrogen receptors were seen in cells in collagen-rich regions and within the interfascicular matrix area. In contrast with western blot results, there were no significant differences observed in ERα staining signal across ages (Figure 6A,B) other than the decreased cellularity in the post-pubescent group and some higher intensity staining in the interfascicular matrix region in the pre-pubescent group (Figure S4). This increased regional staining difference was also seen for GPR30 primarily in the pre-pubescent group. Across age, there was decreased staining intensity for GPR30 in the post-pubescent group, similar to the differences in gene (*GPER1*) and protein (GPR30) concentration (Figure 6C,D). Positive staining was observed for all animals, except for one post-pubescent animal that had a lack of positive ERα staining (Figure 6E). A similar staining pattern was observed across age for the PL bundles (Figure S5). Positive staining for ERα and GPR30 was validated with positive uterine tissues (Figure S6).

**Figure 6.**
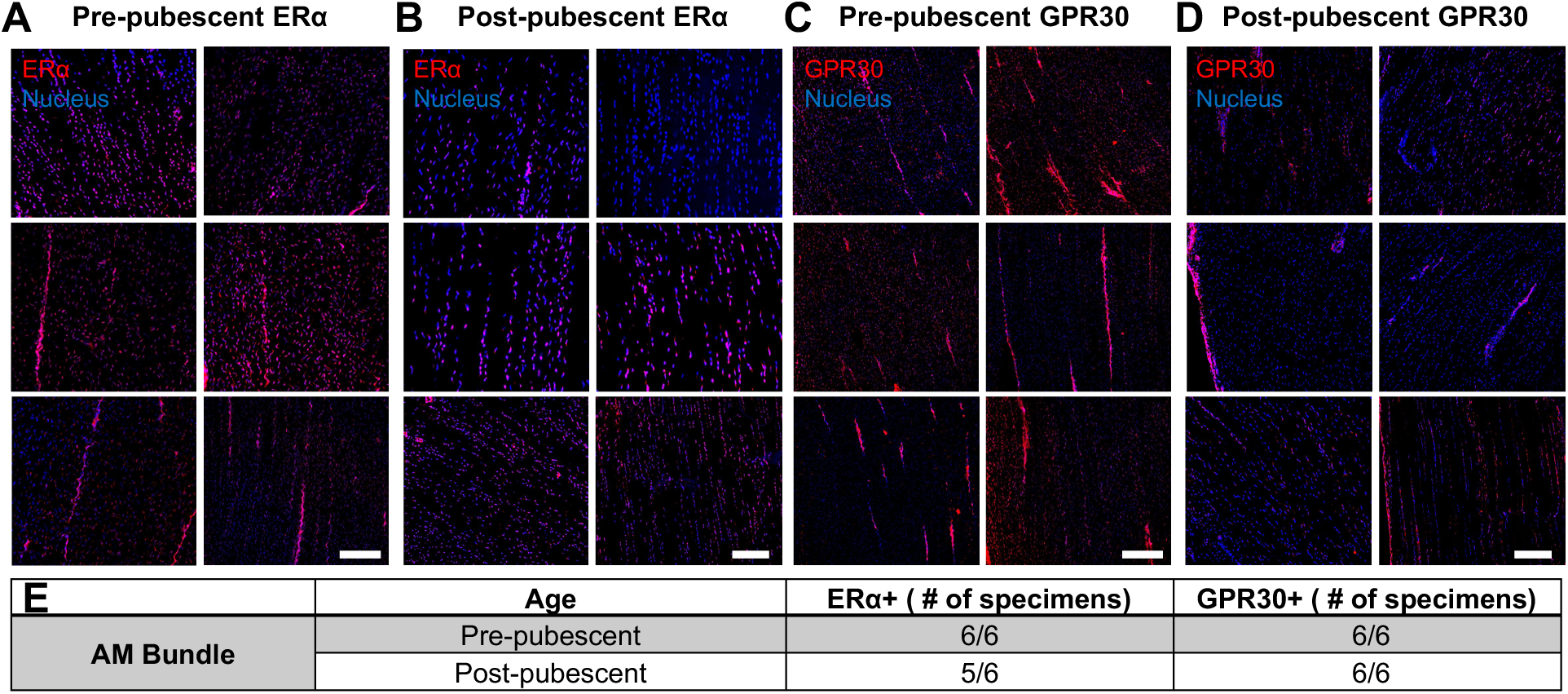
Spatial expression of ERα and GPR30 across age in the AM bundle from immunofluorescence imaging. ERα immunostaining of the AM bundle for all six specimens at (A) pre-pubescence and (B) post-pubescence. GPR30 immunostaining for all six specimens at (C) pre-pubescence and (D) post-pubescence. (E) Total specimens staining positive for either ERα or GPR30. Scale bars 100 µm.

### Changes in ACL Organization Throughout Growth

Histologically, there were distinct differences between pre- and post-pubescent tissues (6/6 pre-pubescent and 6/6 post-pubescent animals analyzed, Figure 7). The cellularity was higher in the pre-pubescent ACLs for each bundle (Figure 7A). The fascicles were also thicker in the post-pubescent group compared to the pre-pubescent ACLs observed in both the picrosirius red and H&E staining. The cellularity and collagen intensity were similar between AM and PL bundles for both age groups, whereas the proteoglycan staining was less intense in the PL bundles compared to the AM bundles (Figure 7B). Comparing between bundles, the PL bundles also had fascicles that were oriented both longitudinally and transversely whereas the AM bundle had fascicles primarily oritented longitudinally, likely from the twisting geometry of the PL bundle. There were no notable histological differences in collagen or proteogylcan distribution across age.

**Figure 7.**
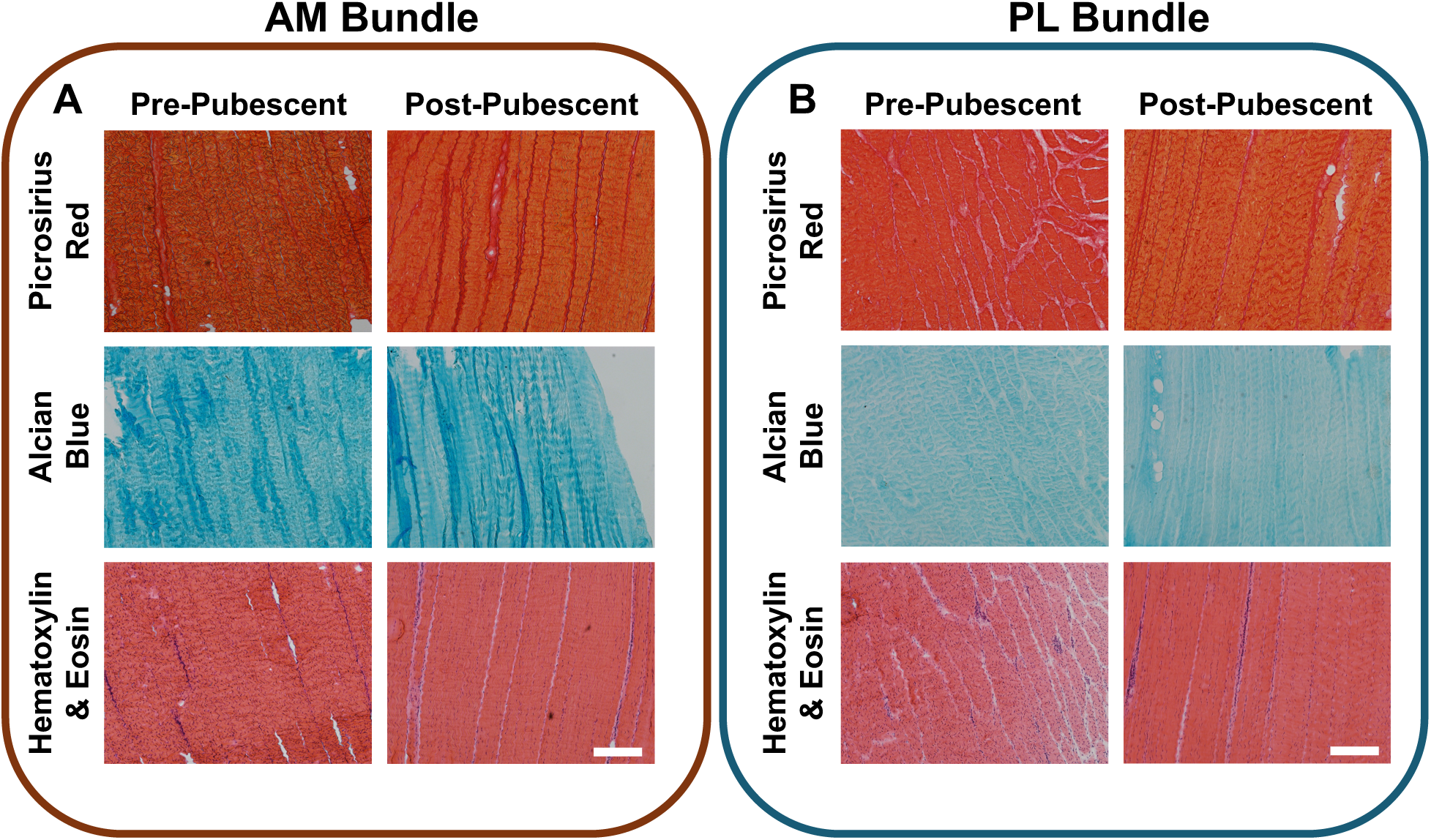
Histological differences in ACL bundle structure across age. Pre- and post-pubescent (A) AM bundles and (B) were stained for collagen organization (picrosirirus red), proteoglycans (alcian blue), and general matrix and nuclei (H&E). Scale bars 100 um.

## Discussion

The findings of this work provide evidence that the sensitivity of the ACL to estradiol may vary across growth, specifically prior to and following puberty. The pre-pubescent AM bundle had greater gene expression of collagen 1, elastin, and tenomodulin compared to the post-pubescent AM bundle, whereas the post-pubescent AM bundle had greater secondary gene expression of collagen 3, lumican, and COMP. In terms of the hormone receptors, the post-pubescent animals had greater gene expression of *ESR1* which encodes for the ERα protein compared to the pre-pubescent group which had minimal levels of *ESR1*. The opposite was true for *GPER1* which encodes for the GPR30 protein, where pre-pubescent AM bundles had greater expression than post-pubescent AM bundles. A similar finding was seen in the protein expression of these two estrogen receptors, though statistical significance was only achieved for ERα. There were minimal differences in gene expression and hormone receptor expression across bundle. These findings confirm our hypothesis that there would be differences in gene expression and hormone receptor expression across age, but not between bundles. From the immunofluorescence, these receptors were seen homogeneously within the post-pubescent and pre-pubescent tissues. Histologically, there were structural and cellularity differences across age as expected.

The transcriptional differences seen across age were in line with expected physiological processes across growth. The higher expression of multiple collagens, including collagen type 1, tenomodulin, and elastin in the pre-pubescent group indicates that the cells within the ACL are focused on matrix synthesis and maturation. Collagen turnover in juvenile and early adolescent human tendons and ligaments is known to be greater than in late adolescence and adulthood ^36^. It was interesting that collagen type 2, a typical marker for cartilaginous tissue, was discovered within the pre-pubescent AM bundle. It is possible these young cells are more complex ligament fibroblasts or cells near the fibrocartilaginous enthesis of the ACL were included. Pre-pubescent AM bundles also have greater *TRPV4* expression that encodes a calcium ion channel important for mechanosensing ^37,38^. This suggests that pre-pubescent AM bundles may be more mechanosensitive than post-pubescent AM bundles, but additional work is needed. In the post-pubescent ACL, there was greater expression of COMP, lumican, BMP12, and collagen 3. Both collagen 3 and BMP12 have been shown to be important for response to injury ^39–41^, indicating that the post-pubescent AM bundle has shifted to more of a homeostatic remodeling state by late adolescence. COMP and lumican have important roles in collagen fibrillogenesis ^42,43^. It could suggest greater turnover of these proteins whereas the expression of primary ligament genes has subsided by this stage of development.

When looking at hormone receptors in the ACL across age, there were clear variations in the gene and protein expression of ERα. While there are several signaling mechanisms for estrogen in the body, the most common and well-understood pathway is through ERα, a nuclear receptor that binds to estrogen, dimerizes, and behaves as a transcription factor after the recruitment of several additional co-factors ^44^. The substantial difference across age signifies that the ACL could be more responsive to circulating levels of estrogen following sexual maturation, which are significantly greater compared to pre-pubescent levels ^45,46^. Estrogen exposure has been shown to have distinct impacts on ACL fibroblasts, including decreasing lysyl oxidase (LOX) activity ^27^, increasing matrix metalloproteinase activity in conjunction with relaxin ^47^, and decreasing collagen synthesis ^26^. A decrease in LOX activity leads to weaker collagen-rich tissues as LOX aids in collagen crosslinking. This paired with an increase in MMP expression which breaks down extracellular matrix proteins and a decrease in collagen synthesis all could contribute to an increased risk of ACL injury in teenage athletes. Changes in ACL responsiveness to estrogen across age may help provide more clarity on the timeline when the ACL begins to be impacted by circulating estrogen. The increase in this receptor expression may also shift with short-term changes in hormone concentrations, yet tracking expression across cycle phase would be necessary to confirm this. Gene expression of *ESR2* was also detected and averaged greater in post-pubescent animals, yet it was variable between animals.

Relative to *ESR1*, an opposite pattern was found for *GPER1* and *PGR*, which had greater gene expression in the pre-pubescent ACLs. The GPR30 protein allows for a rapid response to estrogen through nongenomic signaling, such as cAMP production, calcium mobilization, and activation of ERK and PI3K pathways ^48,49^. These rapid responses may be responsible for acute changes in the ACL throughout the reproductive cycle. Non-genomic estrogen signaling through GPR30 has been implicated in rapid cytoskeletal re-organization of female dermal fibroblasts ^50^, cardiac fibroblast proliferation suppression ^51^, and cancer associated fibroblast migration promotion ^52^. How these signaling mechanisms relate to larger scale structural and mechanical changes in the ACL, however, is still unknown, especially when estrogen levels are low pre-puberty. Similar to ERα, GPR30 had off target molecular weights on the western blot in a similar molecular weight ∼130 kDa as seen for ERα, indicating that perhaps there is some off-target binding of the antibodies. *PGR* expression also had greater gene expression in the pre-pubescent ACLs, but even less is known about the impact of progesterone prior to puberty.

Spatially, both ERα and GPR30 are expressed homogenously throughout the tissue where ERα primarily localized with the nucleus and GPR30 localized to the cytosol and membrane. This is in alignment with previous data showing sex hormone receptors in the human ACL ^24^, indicating potential similar mechanisms at play between the pig ACL and the human ACL. The expression pattern suggests that local estrogen concentrations may impact fibroblasts, synoviocytes, endothelial cells, and other immune cells populations within the ACL. Interestingly, pre-pubescent tissues also had positive estrogen receptor staining despite the low concentration detected on the western blot which might indicate that the higher molecular weights seen in the total western blot may be estrogen receptor protein complexes. Previous immunohistochemistry data showed progesterone receptors in the adult ACL, yet progesterone receptor quantification in the samples in this work was not possible because there was minimal antibody binding to the samples or to the positive uterine control. Together, these results highlight the value of western blotting and the limitations of relying on immunofluorescence staining to assess protein expression. Progesterone receptor expression will need to be confirmed through future work and a validated antibody.

When comparing between AM and PL bundles, there were minimal transcriptional differences at either age. There was only one gene determined to have significantly greater expression in the PL bundle compared to the AM bundle for pre-pubescent samples (*ADAMTS4*). However, qualitatively, there does appear to be greater differences between the AM and PL bundles specifically in pre-pubescent ACL tissues, compared to differences in post-pubescent tissues. Perhaps, this points to the greater cellular variation in the AM and PL bundles at pre-pubescent ages. For hormone receptors, there were no differences between AM or PL bundles for either age groups. This is matched in the protein expression of ERα and GPR30. Similarly, there are no spatial distribution differences between AM and PL bundle receptor expression. While there are clear structural and cellularity differences across growth, there are minor differences between bundles other than some off-axis fascicles in the PL bundle that are likely from the twisting geometry of the PL bundle ^53^. These results suggest minor or undetectable differences between the AM and PL bundles across age.

This study is limited by the small sample size and the lack of control over the cycling nature of the post-pubescent animals. Serum measurements and gross observation of the ovaries in a subset of animals provided more insight into potential differences in estrous cycle phase for the post-pubescent animals. Three animals that had serum analyzed had estradiol values greater than 20 pg/ml, which indicates that they are likely in active estrus or near estrus (either immediately prior to or following ovulation) based on previously reported values of the estradiol spike prior to estrus ^54,55^. One of the four pigs had low estradiol and low progesterone based on the spike of progesterone in pigs previously reported to be between 20 – 30 ng/ml, indicating that it was likely days prior to estrus or days after estrus before progesterone increases ^54,55^. Active estrus detection was not monitored behaviorally in any animal and only one serum sample was taken for a subset of animals; thus the true estrus phase was not definitively known. Further work seeking to make conclusions about transcriptional or biological variations in the ACL across the reproductive cycle of the pig should include longitudinal serum measurements and behavioral and physiological observations. The specimens used in this study also came from multiple research studies, so the exact history of the animal and the potential impact on the developing knee joint was not known. Another limitation was the small scope of the gene expression panel used, which only had 45 preselected genes related to tendon and ligament phenotypes and hormone receptors. The results analyzed also only came from female animals instead of males and females. To better understand a more comprehensive transcriptional and/or proteomic landscape across age and sex, more thorough tools like RNAseq and proteomic analysis should be utilized. Finally, although we found statistically significant differences across age, we did not know what effect sizes to expect. Thus, it is possible that the study was underpowered for smaller effect sizes (for example, data in Fig. 4D and 5D) and that larger samples would find statistically significant differences. Despite these limitations, this study lays the groundwork for distinct differences in the matrix-related and hormone receptor-related differences.

Overall, the structural and cellularity shifts in the developing ACL across adolescence are matched by changes in the cellular gene expression where there is an initial focus on extracellular synthesis and maturation that moves toward a homeostatic focus. Shifts are also seen in the hormone receptor profiles, with greater ERα gene expression post-puberty and greater GPR30 gene expression pre-puberty. These differences further support the idea that the ACL is hormonally responsive and that there are multiple estrogen signaling pathways utilized in the adolescent ACL.

While a pig model was used in this study, the matching hormone receptor appearance in the human and pig ACL indicates similar potential mechanisms of impact between the two species. The findings here motivate further work to understand more specifically what changes sex hormones initiate within the ACL that may help explain the increased injury risk in adolescent females.

## Acknowledgments

We would like to thank Laboratory Animal Resources (NC State) and the Clinical Endocrinology Lab (UC Davis) for their contributions to this work. The authors acknowledge the use of Claude Sonnet 4.5 (accessed February 2026) to support creation of the abstract. All AI-generated suggestions were reviewed, revised, and approved by the authors, who take full responsibility for the accuracy and integrity of the work.

## Funding Data

- Division of Graduate Education (Grant No. DGE-1746939; Funder ID: 10.13039/100000082).
- National Institute of Arthritis and Musculoskeletal and Skin Diseases (Grant No. 2R01AR071985; Funder ID: 10.13039/100000069. The content is solely the responsibility of the authors and does not necessarily represent the official views of the National Institutes of Health.

