## Supplemental Data for "Estrogen Receptor Expression Changes After Puberty in the Porcine Anterior Cruciate Ligament"

**Supplemental Material**

**Table S1.** Summary table of all genes analyzed in the Nanostring custom porcine codeset grouped by category.

| **Gene Category** | ***Gene Symbol*** |
|---|---|
| Extracellular Matrix | *ACAN, BGN, COL1A1/2, COL2A1, COL3A1, COL5A1, COL6A1, COMP, DCN, ELN, FMOD, FN, LUM, PRG4, VCAN* |
| Tendon & Ligament | *MKX, SCX, TNC, TNMD* |
| Matrix Remodeling & Sensing | *ACTA2, ADAMTS2/4/5, LOX, LOXL2, MMP1/13/3/9, PIEZO1, TRPV4* |
| Hormone Receptors | *ESR1/2, GPER1, PGR* |
| Growth Factors | *FGF2, GDF7/BMP12, TGFB1/2/3, VEGF* |
| Housekeeping | *ACTB, GAPDH, HPRT1* |

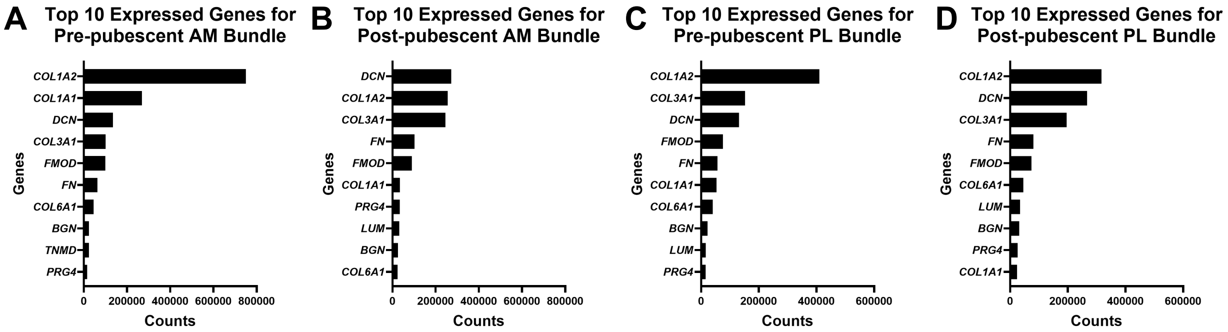


**Figure S1.** Top expressed genes based on raw count for all bundles and ages tested. Raw counts for the top 10 genes expressed of (A) pre-pubescent AM bundles, (B) post-pubescent AM bundles, (C) pre-pubescent PL bundles, and (D) post-pubescent PL bundles. Data expressed as mean counts.

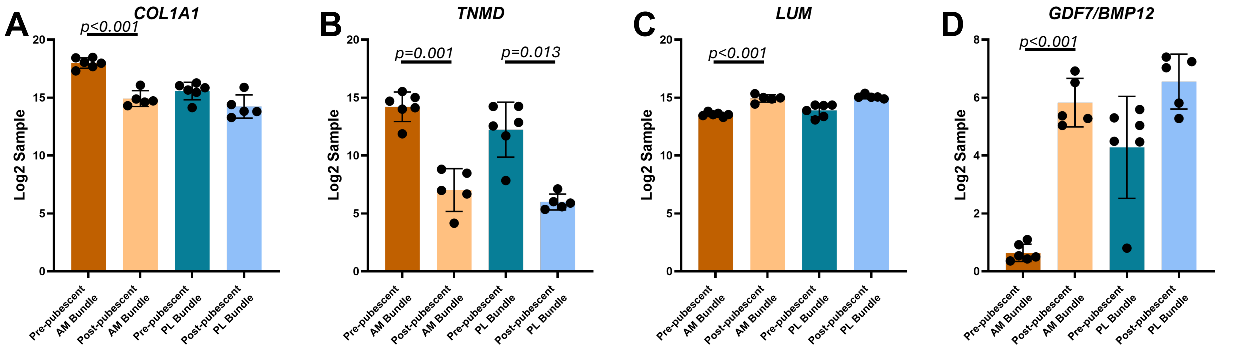


**Figure S2.** Extracellular matrix and growth factor gene expression differences across age. Log2 of raw counts between bundle and age for (A) *COL1A1*, (B) *TNMD*, (C) *LUM*, and (D) *GDF7/BMP12*. Individual data points represent biological specimens and data expressed as mean ± s.d. Bars indicate $p<0.05$.

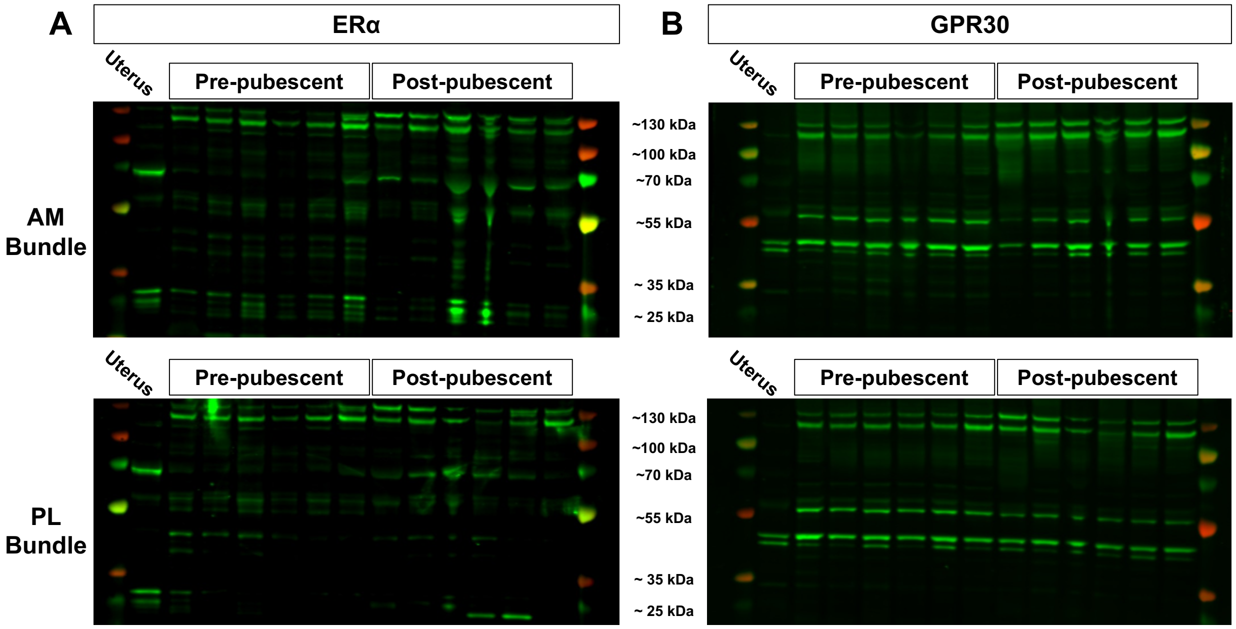

**Figure S3.** Complete membrane immunoblotting of ERα and GPR30 across age and bundle with positive uterus loaded control. Positive ERα bands at expected ~66 kDa and at high (~130 kDa) and low (~25 kDa) molecular weights for (A) AM and PL bundles. Positive GPR30 bands at expected ~42 kDa and at high (~58 kDa and ~130 kDa) molecular weights for (B) AM and PL bundles.

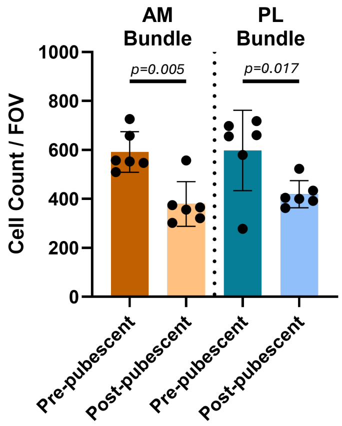


**Figure S4**. Variations in cellularity in ACL tissue across age and bundle. Cells counted within the field of view from immunofluorescence image utilizing the analyze particles function in ImageJ. Individual data points represent biological specimens and data expressed as mean ± s.d. Bars indicate $p<0.05$.

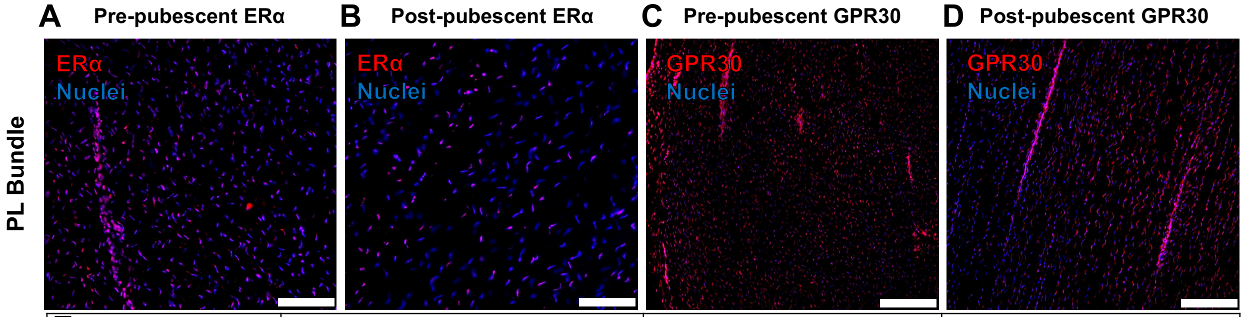


| E | Age | ERα+ ( # of specimens) | GPR30+ ( # of specimens) |
|---|---|---|---|
| PL Bundle | Pre-pubescent | 6/6 | 6/6 |
| | Post-pubescent | 6/6 | 6/6 |

**Figure S5.** Spatial expression of ERα and GPR30 across age in the PL bundle from immunofluorescence imaging. Representative ERα immunostaining of the PL bundle at (A) pre-pubescence and (B) post-pubescence. Representative GPR30 immunostaining of the PL bundle at (C) pre-pubescence and (D) post-pubescence. (E) Total specimens staining positive for either ERα or GPR30. Scale bars 100 μm.

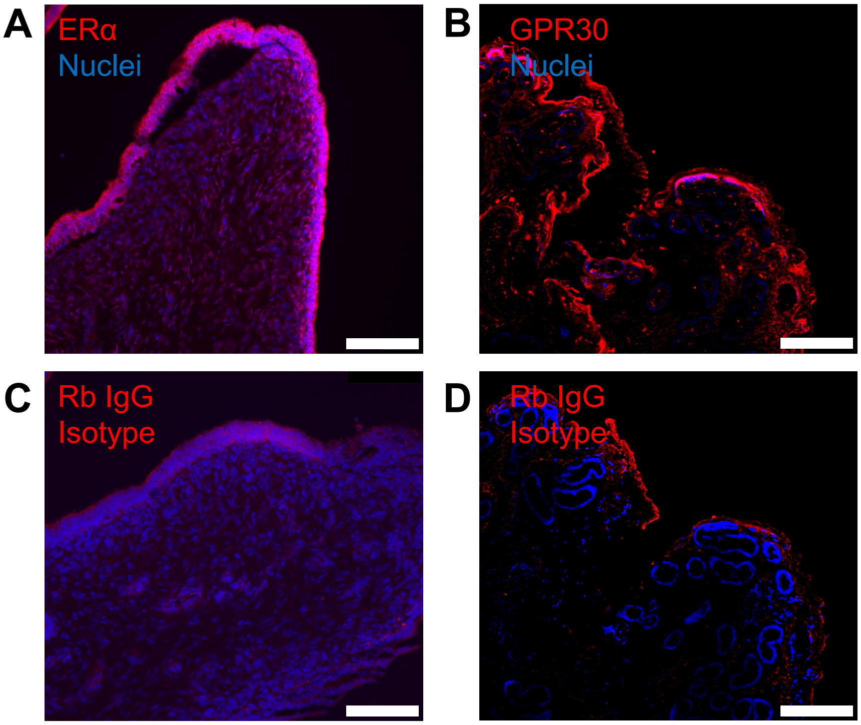


**Figure S6.** Hormone receptor antibody validation in porcine uterus tissue (positive control). Porcine uterus staining for (A) ERα and (B) GPR30. (C,D) Corresponding rabbit IgG isotype control staining in the porcine uterus. Scale bars 100 μm.
